# Three-way clustering of multi-tissue multi-individual gene expression data using constrained tensor decomposition

**DOI:** 10.1101/229245

**Authors:** Miaoyan Wang, Jonathan Fischer, Yun S. Song

## Abstract

The advent of next generation sequencing methods has led to an increasing availability of large, multi-tissue datasets which contain gene expression measurements across different tissues and individuals. In this setting, variation in expression levels arises due to contributions specific to genes, tissues, individuals, and interactions thereof. Classical clustering methods are ill-suited to explore these three-way interactions, and struggle to fully extract the insights into transcriptome complexity and regulation contained in the data. Thus, to exploit the multi-mode structure of the data, new methods are required. To this end, we propose a new method, called *MultiCluster*, based on constrained tensor decomposition which permits the investigation of transcriptome variation across individuals and tissues simultaneously. Through simulation and application to the GTEx RNA-seq data, we show that our tensor decomposition identifies three-way clusters with higher accuracy, while being 11x faster, than the competing Bayesian method. For several age-, race-, or gender-related genes, the tensor projection approach achieves increased significance over single-tissue analysis by two orders of magnitude. Our analysis finds gene modules consistent with existing knowledge while further detecting novel candidate genes exhibiting either tissue-, individual-, or tissue-by-individual specificity. These identified genes and gene modules offer bases for future study, and the uncovered multi-way specificities provide a finer, more nuanced snapshot of transcriptome variation than previously possible.

## Introduction

Owing to advances in high-throughput sequencing technology, multi-tissue expression studies have provided unprecedented opportunities to investigate transcriptome variation across tissues and individuals (Lonsdale et al. 2013; Melé et al. 2015; Walker et al. 2004; Nica et al. 2011; Hawrylycz et al. 2012). A typical multi-tissue experiment collects gene expression profiles (e.g. via RNA-seq or microarrays) from different individuals across a number of different tissues, and variation in expression levels often results from complex interactions among the different modes of the data (Melé et al. 2015). For example, a group of genes may perform coordinated biological functions in certain contexts (e.g. specific tissues or individuals), but may behave differently in other settings through tissue- and/or individual-dependent gene regulation mechanisms. An improved understanding of these interactions will yield insight into fundamental biological questions with clinical applications such as changes in cellular function (Dönertaş et al. 2017), personalized transcriptomics (Montgomery and Dermitzakis 2011), and disease susceptibility (Melé et al. 2015).

Clustering has proven useful to reveal latent structure in high-dimensional expression data (Kluger et al. 2003; Kiselev et al. 2017; Wiwie et al. 2015). Traditional clustering methods (such as K-means, PCA, and t-SNE (Maaten and Hinton 2008)) assume that gene expression patterns persist across one of the different contexts (either the tissue or individual mode), or assume that samples within one mode are i.i.d. or homogeneous. Direct application of these algorithms to multi-tissue expression data requires concatenating all available samples from different tissues into a single matrix, precluding potential insights into tissue × individual specificity (Bahcall 2015). Alternatively, inferring gene modules separately for each tissue ignores commonalities among tissues and may hinder the discovery of differentially expressed genes that characterize tissues or tissue groups. Likewise, individuals vary by their biological attributes (such as ethnicity, gender and age), and ignoring such heterogeneity impedes the accurate estimation of gene- and/or tissue-wise correlation (McCall et al. 2016). The development of a statistical method that integrates multiple modes simultaneously is therefore essential for elucidating the complex biological interactions present in multi-tissue multi-individual gene expression data.

Several methods have been proposed in multi-tissue multi-individual expression studies, but they are often unable to fully exploit the three-mode structure of the data. Pierson et al. (2015) propose a hierarchical transfer learning algorithm to learn gene networks in which they first construct a global tissue hierarchy based on mean expression values and subsequently infer gene networks for each tissue conditioned on the tissue hierarchy. Dey et al. (2017) instead use topic models to cluster samples (i.e. tissues or individuals), and based on tissue clusters they identify genes that are distinctively expressed in each cluster. Both algorithms take a two-step procedure to uncover expression patterns in tissues and genes. Other methods aim to take a one-shot approach by identifying subsets of correlated genes that are exclusive to, for example, female individuals. Gao et al. (2016) adopt the biclustering framework and propose decomposing the expression matrix into biclusters consisting of subsets of samples and features with latent structure unique to the overlap of particular subsets. However, in the case of multi-tissue measurements across individuals, concatenating the data sample-wise to create a single expression matrix will not explore the threeway interactions among genes, tissues, and individuals. A more recent work (Hore et al. 2016) develops sparse decomposition of arrays (SDA) for multi-tissue expression experiments. Because their focus is not on clustering tissues or individuals, the proposed i.i.d. prior on individual/tissue loadings may not be powerful enough to detect tissue- and individual-wise correlation.

In this paper, motivated by recent success in tensor decomposition (Wang and Song 2017; Kuleshov et al. 2015; Sidiropoulos et al. 2017), we address the aforementioned challenges by using a tensor-based approach to jointly cluster genes, tissues, and individuals. We utilize triplets of sorted loading vectors in a constrained tensor decomposition to represent three-way clusters specific to subsets of genes, tissues, and individuals. This approach handles heterogeneity in each mode and automatically learns the interplay among different modes of the data. We propose a simple but principled statistical procedure to characterize the identified three-way clusters on the basis of available metadata. Our method uncovers several different types of gene expression modules, including (i) global, shared expression modules; (ii) expression modules specific to certain subsets of tissues; (iii) modules with differentially expressed (DE) genes across individual-level covariates (e.g., age, sex or race); and (iv) expression modules that are specific to both tissues and individuals.

## Results

### Overview of our *MultiCluster* method

Below, we briefly describe a new clustering method, *MultiCluster*, for identifying three-way clustering patterns in multi-tissue multi-individual gene expression data. Additional details can be found in Materials and Methods.

As illustrated in Figure 1a, multi-tissue multi-individual gene expression measurements can be organized into a 3-way array, or order-3 tensor, with gene, tissue, and individual modes. Our goal is to identify subsets of genes that are similarly expressed in subsets of tissues and individuals; mathematically, this reduces to detecting 3-way blocks in the expression tensor (Figure 1b). These local blocks may correspond to, for example, gene expression modules that are active in some but not all tissues and individuals. We utilize the flexible tensor decomposition framework to directly identify gene modules in a tissue × individual specific fashion, which traditional clustering methods would fail to capture.

**Figure 1.**
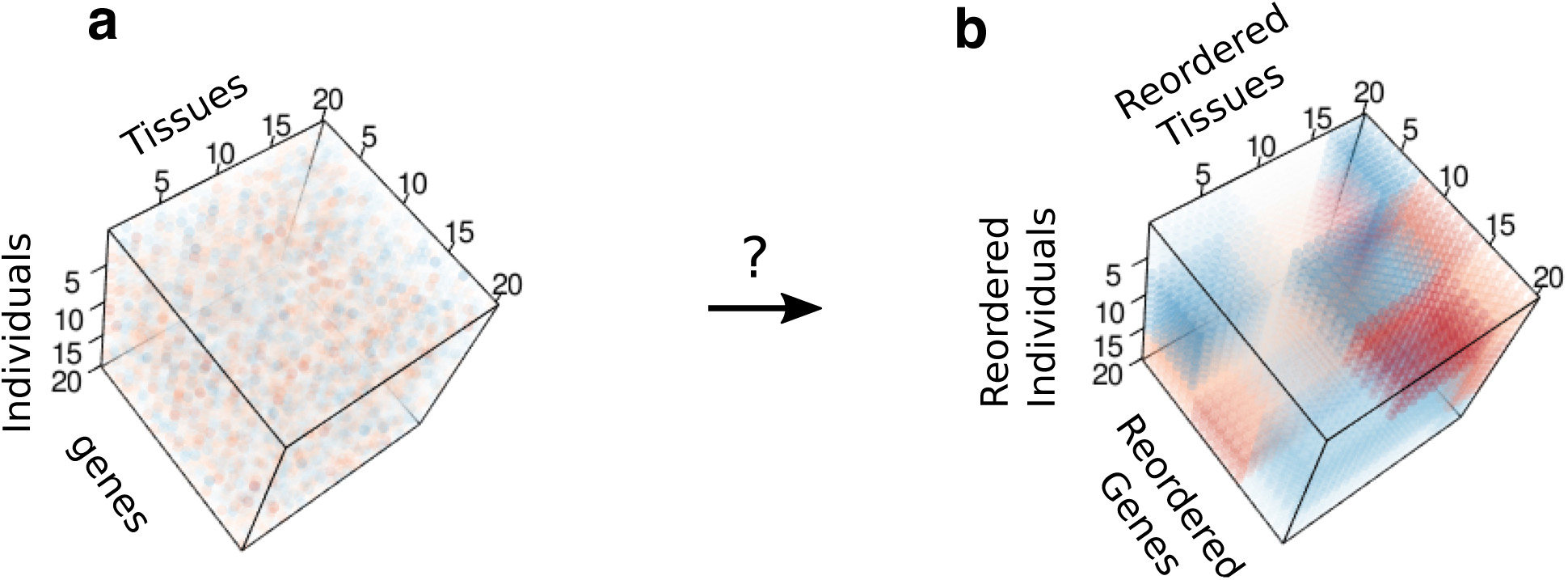
Three-way clustering problem. (a) Input tensor of gene expression. (b) Shuffled, de-noised output tensor containing local blocks. Both (a) and (b) are color images of a data tensor 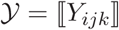, with each entry colored according to the value of *Y_jk_*.

Figure 2a-d provide a schematic illustration of the *MultiCluster* method. Briefly, *MultiCluster* takes as input the multi-tissue multi-individual gene expression data, 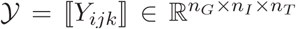, where *Y_ijk_* represents the expression value (possibly after a suitable transformation) of gene *i* measured in individual *j* and tissue *k* (Figure 2b). Our algorithm decomposes *Y* into a sum of rank-1 components,

**Figure 2.**
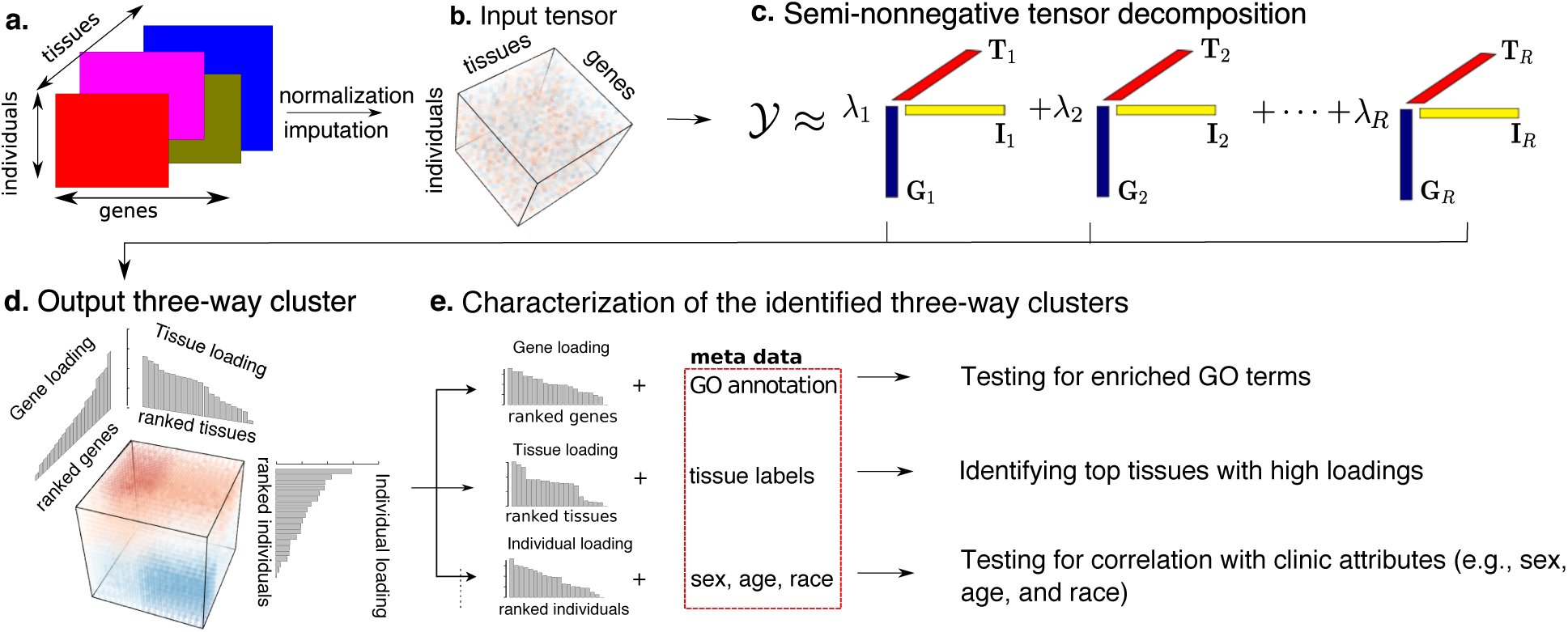
Schematic diagram of *MultiCluster* algorithm. (a) Multi-tissue gene expression data. (b) Input expression tensor after normalization and imputation. (c) The algorithm decomposes the expression tensor into a set of rank-1 tensors, ***G_r_*** ⊗ ***I_r_*** ⊗ ***T_r_***, where ***G_r_***, ***I_r_***, and ***T_r_*** are, respectively, gene, individual, and tissue singular vectors. (d) Each three-way cluster is represented by the three sorted singular vectors. (e) We utilize metadata, such as GO annotation, tissue labels, and individual-level covariates, to identify the sources of variation in each loading vector.

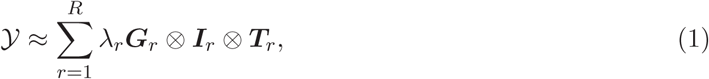

where λ_1_ ≥ λ_2_ ≥ … ≥ λ_*R*_ ≥ 0 are singular values in descending order, and ***G**_r_, **I**_r_* and ***T**_r_* are norm-1 loading vectors indicating the relative contribution of each gene, individual, and tissue to the *r*-th component (Figure 2c). We interpret the rank-1 tensor ***G**_r_* ⊗ ***I**_r_* ⊗ ***T**_r_* as the basic unit of expression pattern (called an expression module), in which the (*i,j,k*)-th entry of ***G**_r_* ⊗ ***I**_r_* ⊗ ***T**_r_* is the multiplicative product of the corresponding entries in the three modes, i.e., (***G**_r_* ⊗ ***I**_r_* ⊗ ***T**_r_*)_(*i,j,k*)_ = *G_r,i_I_r,j_T_r,k_*. Each module may represent a particular interaction of biological processes. Genes with large *G_r_*-values are affected more greatly by the tissues and individuals in the module *r*, whereas these effects are also greater for individuals with larger *I_r_*-values and tissues with larger *T_r_*-value (Figure 2d). To facilitate interpretation, we impose entrywise non-negativity on the tissue loading vectors *T_r_* by zeroing negative values of *T_r,k_*. By examining the entries in the tissue vectors *T_r_*, one can learn the tissue activity pattern of the associated module. Note that no sign constraint is imposed on individual and gene loadings, therefore the method handles mixed-sign data tensors. We adopted our earlier algorithm (Wang and Song 2017) based on successive rank-1 approximations to solve the tensor decomposition (1).

Extending earlier work (Alter et al. 2000; Omberg et al. 2007), we refer to the loading vectors ***G**_r_*, ***I**_r_*, ***T**_r_* as “eigen-genes”, “eigen-individuals”, and “eigen-tissues”, respectively. To characterize the biological significance of the inferred modules, we utilize metadata to identify the sources of variation in each loading (Figure 2d). For each eigen-gene, we identify gene ontology (GO) terms enriched among the top-ranked genes, and annotate each eigen-tissue using its positively-loaded tissues. An expression module is declared “tissue-specific” if the eigen-tissue is dominated by only a few tissues, and “age-, sex-, or race-related” if the eigen-individual loadings are correlated with age, sex, or race, respectively. We also test whether these individual-level covariates can account for the variation in eigen-individuals and employ tensor projection onto the eigen-tissues to pinpoint the genes which drive the signal in covariate-associated expression modules.

### Summary of simulation results

To test the performance of *MultiCluster*, we carried out a simulation study and compared the results with a number of alternative methods. The simulations served two purposes: 1) to evaluate the ability of *MultiCluster* to recover multi-way clusters under various cluster models, and 2) to assess the power of our tensor-based procedure to detect DE genes and compare with the power in standard single-tissue analysis.

*Accuracy of three-way clustering*. We applied *MultiCluster* to identify 3-way blocks in noisy expression tensors simulated from three different cluster models: additive-, multiplicative-, and combinatorial-mean models, as detailed in Materials and Methods. We compared the recovery accuracy with two recently developed tensor methods: (i) sparse decomposition of arrays (*SDA*) (Hore et al. 2016), and (ii) tensor higher-order singular value decomposition (*HOSVD*) (Omberg et al. 2007). Both *MultiCluster* and *SDA* are built upon the Canonical Polyadic (CP) decomposition which decompose a tensor into a sum of rank-1 matrices, whereas *HOSVD* decomposes a tensor into a core tensor multiplied by an orthogonal matrix in each mode.

As seen in the example tensors depicted in Figure 3, **MultiCluster** is able to recover the block structure well in all three scenarios, demonstrating its robustness to model misspecification. We used the relative estimation error (up to permutation) across 50 simulations to assess the recovery accuracy of each method (Materials and Methods). We found that **MultiCluster** consistently outperforms the other two methods (Figure 3a-c). For the additive and multiplicative models, the two non-Bayesian methods (**MultiCluster** and *HOSVD*) tend to recover the block structure better than SDA (Figure 3a-b). A possible explanation is that *SDA* is designed to cluster genes rather than tissues and individuals, so the i.i.d. prior imposed on tissues/individuals may not be powerful enough to detect local blocks, especially the blocks are small. It may also be due the algorithmic stability of *MultiCluster* relative to *SDA*; the latter usually requires multiple restarts in order to reduce spurious components (Hore et al. 2016). For the more complicated combinatorial model, however, the two CP decompositions (*MultiCluster* and *SDA*) yield lower errors than the *HOSVD* (Figure 3c). Note that neither *MultiCluster* nor *SDA* forces orthogonality in the loading vectors; instead, they adopt some sparsity and regularity (tissue nonnegativity for *MultiCluster* and gene sparsity for *SDA*). The way we incorporated tissue non-negativity essentially introduces zeros into the rank-1 tensor output, thereby fitting the 3-way blocks more flexibly in a local fashion.

**Figure 3.**
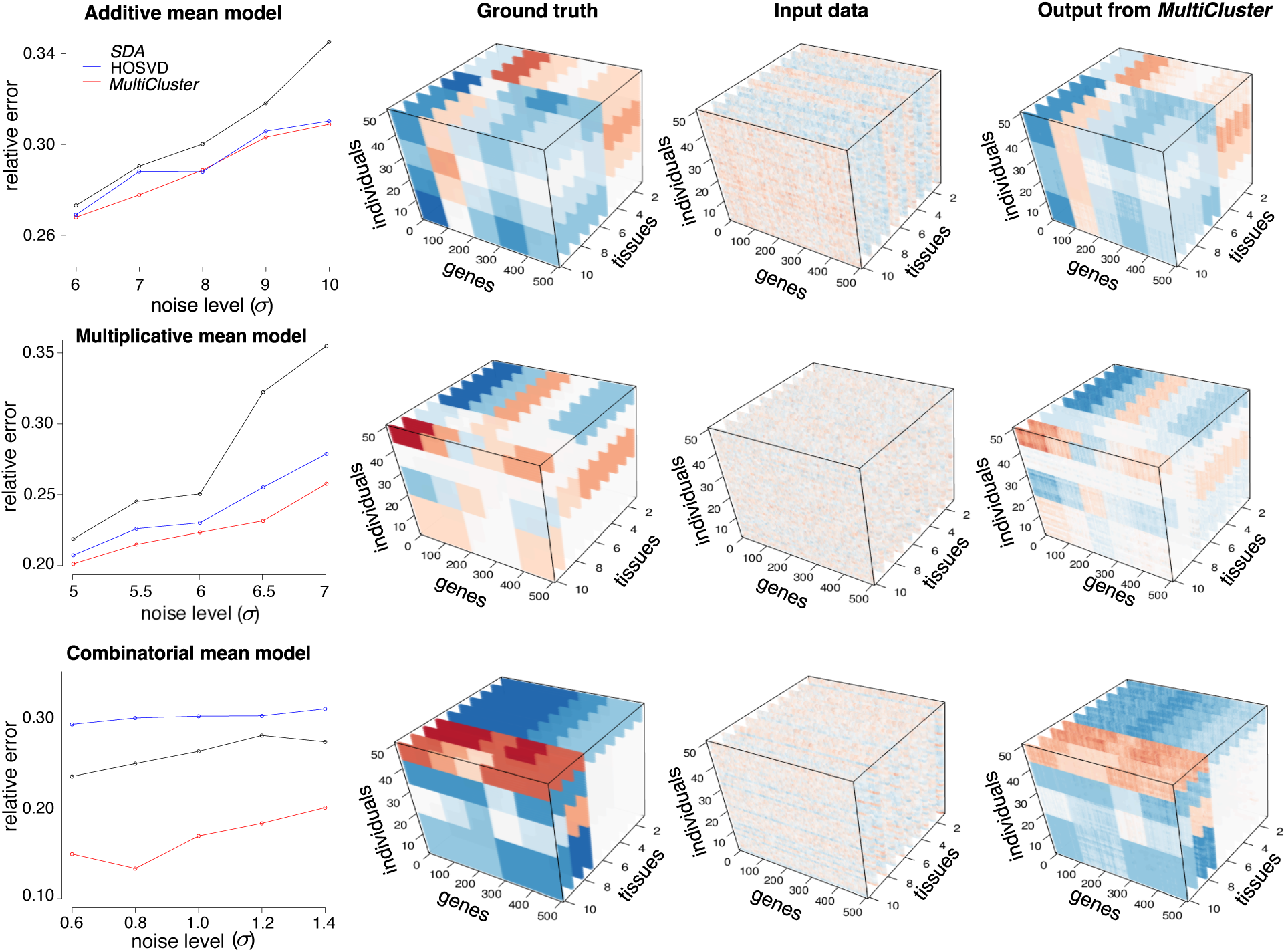
Three-way clustering performance in simulations. The first column shows the recovery accuracy of different tensor-based methods. The columns 2–4 are color images of example tensors in simulations under different block-mean models. The example tensor 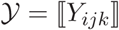 1 consists of 500 (genes) × 50 (individuals) ×10 (tissues), with each entry colored according to the value of *Y_ijk_*.

*Power to detect differentially-expressed genes*. To study how our tensor projection procedure affects the ability to detect covariate-associated genes, we simulated age-related genes. Expression tensors were generated according to the additive model with 10 tissues (3 tissue groups), and 100 genes (out of the 500 in total) were randomly chosen to be age-related. Each age-related gene was active in at least one tissue cluster, and for every tissue in the active cluster, the age effect size was independently drawn from a Unif[0, 0.05] (up-regulated) or Unif[–0.05,0] (down-regulated) distribution. We decomposed each simulated tensor into *R* = 3, 5, 10 components and applied our tensor-projection procedure (Materials and Methods) to test for age-related genes. We declared a gene age-related if its *p*-value is less than the nominal significance level in at least one of the *R* eigen-tissues. To see whether we have improved power over single-tissue tests, we performed standard linear regressions in each tissue separately and declared a gene age-related if its *p*-value is less than the nominal level in at least one of the 10 tissues.

Figure 4 shows the receiver operating characteristic (ROC) curves that compare the true positive rate vs. false positive rate for each method. We found that the testing procedures based on tensor projection have higher detection power than the single-tissue analysis, demonstrating the advantage of tensor-based methods in incorporating information across similar tissues. In particular, the power seems to be stable when the decomposition rank *R* is increased from 3 (the number of latent tissue groups) to 10 (the number of total tissues). It is worth noting that the number of association tests conducted in the tensor projection based approaches are fractions of those conducted when tissues are considered one at a time. Since the burden imposed by multiple testing corrections is a common concern in genome-scale testing, our tensor approach is especially attractive because one may perform fewer tests while achieving higher power.

**Figure 4.**
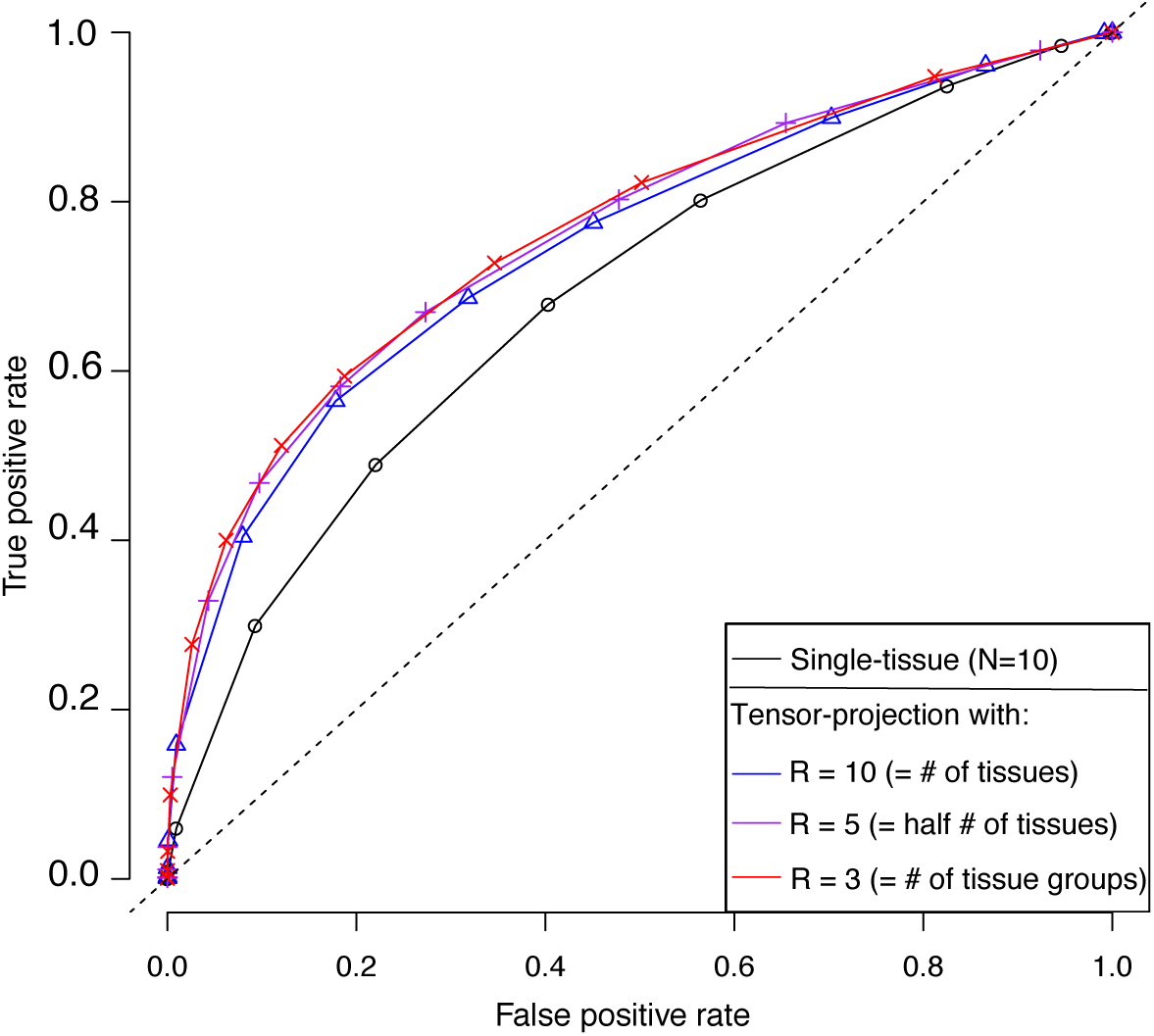
ROC curves for tensor projection and single-tissue analysis. We performed tensor projection and single-tissue analyses to detect age-related genes in simulated expression tensors. For tensor projection, we decomposed each tensor into *R* components with *R* = 3, 5 and 10, and declared an age-related gene if its *p*-value was less than the nominal significance level in at least one of the *R* eigen-tissues. For single-tissue analyses, we tested age-related genes in each tissue separately and declared an age-related gene if its *p*-value was less than the significance level in at least of the 10 tissues. The ROC curves were obtained under various nominal significance levels using 50 simulations.

*Run time*. To compare computational performance, we simulated a large order-3 tensor of 18,000 genes × 500 individuals × 40 individuals. The size of this tensor is to mimic the size of GTEx RNA-seq dataset. We recorded the run times for each method while decomposing this tensor into 10 components. It took 6,047 seconds (≈ 1.7 hrs) for *MultiCluster* and 73,989 seconds (≈ 20.1 hrs) for *SDA* to finish the run. The run time of *HOSVD* (5,849 seconds) is roughly the same as that of *MultiCluster*.

### Identifying expression modules across tissues and individuals in GTEx data

We applied our method to the GTEx V6 gene expression data, which consists of 8,555 RNA-seq samples collected from 544 human individuals across 53 tissues. Each RNA-seq sample measures the read counts of 56,238 annotated transcripts, of which roughly 25,000 are putative protein-coding regions, 10,000 are long non-coding RNAs (lncRNA), and 10,000 are pseudogenes. For convenience, we henceforth refer to all of these transcripts as “genes.” After performing our data processing procedure and removing low-expressed genes (Materials and Methods), we organized the expression measurements into a gene × individual × tissue tensor 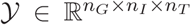, where *n_G_* = 18,481, *n_I_* = 544 and *n_T_* = 44. This tensor, which we named the GTEx tensor, is used in the global tissue analysis.

To investigate the dominant features in the human transcriptome, we performed a global, alltissue clustering analysis to identify gene × tissue × individual expression modules. We applied *MultiCluster* to the GTEx tensor, excluding chromosome Y-linked genes and sex-specific tissues. Supplemental Table S1 summarizes the top expression modules along with their characterization.

*Component I - shared, global expression*. Examining the eigen-tissue loadings informs us in which tissues the corresponding module is active. As expected, the first eigen-tissue and eigen-individual are essentially flat (Supplemental Figure S1a and Supplemental Figure S1c), so this expression module captures baseline global expression common to all samples. The top genes in the corresponding eigen-gene (Supplemental Figure S1b) are mainly mitochondrial genes (15/20 top genes), comporting with their pervasive transcription across tissues as a result of the mitochondrion’s critical role in cellular energy production (Kelly et al. 2012; Melé et al. 2015). In addition, we detected several non-mitochondrial genes, most of which are related to essential protein synthesis functions and eukaryotic cell activities (Supplemental Figure S1d). For example, ACTB encodes highly conserved proteins (Valente et al. 2009) and is known to be involved in various types of cell motility (Fishilevich et al. 2016). Two other nuclear genes, *EEF1A1* and *EEF2*, encode eukaryotic translation elongation factors, and their isoforms are widely expressed in the brain, placenta, liver, kidney, pancreas, heart, and skeletal muscle (Fishilevich et al. 2016).

*Component II - brain tissues*. The second eigen-tissue clearly separates brain tissues from nonbrain tissues, with the pituitary gland being the only non-brain tissue in the cluster (Figure 5a). We note that while not explicitly labeled as a brain tissue, the pituitary gland protrudes from the base of the brain. The sharp decline in tissue loadings (Figure 5a) highlights the distinctive expression pattern in the brain. We found that, in the eigen-individual (Figure 5c), age explains more variation (24.4%, *p* < 2 × 10^−16^) than sex (0.3%, *p* = 0.12) or race (4.3%, *p* = 2.3 × 10^−8^) (Figure 5e; Materials and Methods). The eigen-gene (Figure 5b) produces a gene clustering that is biologically coherent with the brain × aging signal. Indeed, we observed an enrichment of genes from the glutamate receptor signaling pathway (e.g. *CACNG7, CACNG3, GRIN11*; *p* = 1.2 × 10^−20^), chemical synaptic transmission (e.g. *SEZ6, GRM5, GRIA2*; *p* = 1.8 × 10^−16^), excitatory postsynaptic potential (e.g. *RIMS2, CHRNB2, RIMS1*; *p* = 2.4 × 10^−16^), and memory (e.g. *SLC24A2, JPH3, SYT4*; *p* = 1.2 × 10^−11^) (Figure 5d). Among the 899 genes in this cluster, we identified 675 age-related genes using tensor-projection (with significance threshold α = 10-3/899 ≈ 10^−7^ via Bonferroni correction; see Materials and Methods), 556 of which exhibit decreased expression with age (Supplementary Data). The association of brain disease and neurological disorders with age is well-documented, and our findings support that aging affects brain tissues in a manner not shared by other tissues. We present further evidence of multi-way clustering in the brain in Fine structures in subtensors of similar tissues.

**Figure 5.**
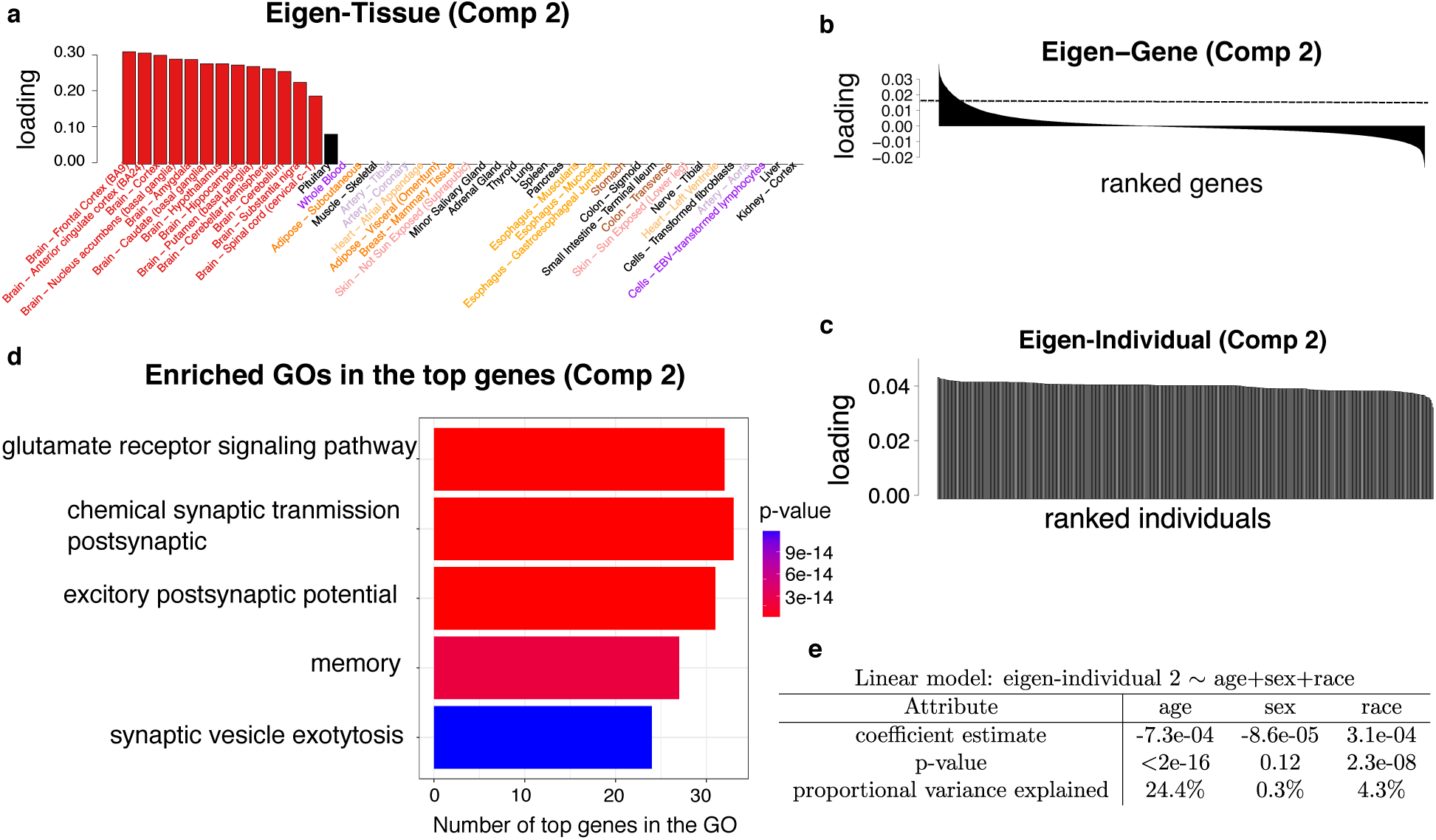
Expression module II - brain tissues. Panels (a)-(c) represent the triplets of sorted singular vectors, whereas panels (d)-(e) represent the biological contexts of the identified threeway cluster. (a) Barplot of the sorted tissue loading vector, where each tissue is colored based on their functional similarity. (b) Barplot of the sorted gene loading vector, where the dotted line represents the threshold for the top genes. (c) Barplot of the sorted individual loading vector. (d) Enriched GO annotations among the top 899 genes identified from the gene loading vector. The enrichment *p*-values are obtained from hypergeometric tests with B-H correction. The GO size on the axis represents the number of top genes that belong to the GO annotation. (e) Linear regression analysis of individual loadings against individual-level covariates (age, sex and race). The proportional variance explained is computed from ANOVA analysis.

*Component III - tissues involved in immune response*. The third component captures an expression module specific to tissues with roles in the immune system. The eigen-tissue is primarily driven by two blood tissues (whole blood and EBV-transformed lymphocytes), the spleen, and the liver (Figure 6a). These tissues serve as mediators of direct immune response (whole blood and lymphocytes), the production and storage of antibodies (spleen), and filtering of antigens (spleen and liver). Our eigen-tissue also reveals the subtle involvement of esophagus in this module (Figure 6a). In fact, mast cells (part of immune system) have been found at the basement membrane and in the lamina propria of normal esophageal mucosa (Collins 2014), reflecting the mixture of cell types in esophagus tissues.

Correspondingly, the eigen-gene loads heavily on immunity-related genes (e.g. *IGHM, FCRL5, IGJ, MS4A1*) (Figure 6b). The eigen-individual does not correlate with any covariate as strikingly as the brain does with age, but we do find a significant correlation with race (explaining 4.5% variation among individuals, *p* = 5.8 × 10^−7^; Figure 6e). The top genes in the eigen-gene are functionally related to the B cell receptor signaling pathway (e.g. *BLK, CD79A, IGHG3*; *p* = 3.0 × 10^−15^), humoral immune response mediated by circulating immunoglobulin (e.g. *IGHM, IGHD, IGHA1*; *p* = 7.5 × 10^−13^), the phagocytosis recognition (e.g. *IGHA2, IGHG1, IGHG2*; *p* = 5.3 × 10^−10^), and plasma membrane invagination (e.g. *AURKB, IGHV3-23, IGHG4*; *p* = 2.1 × 10^−9^) (Figure 6d).

**Figure 6.**
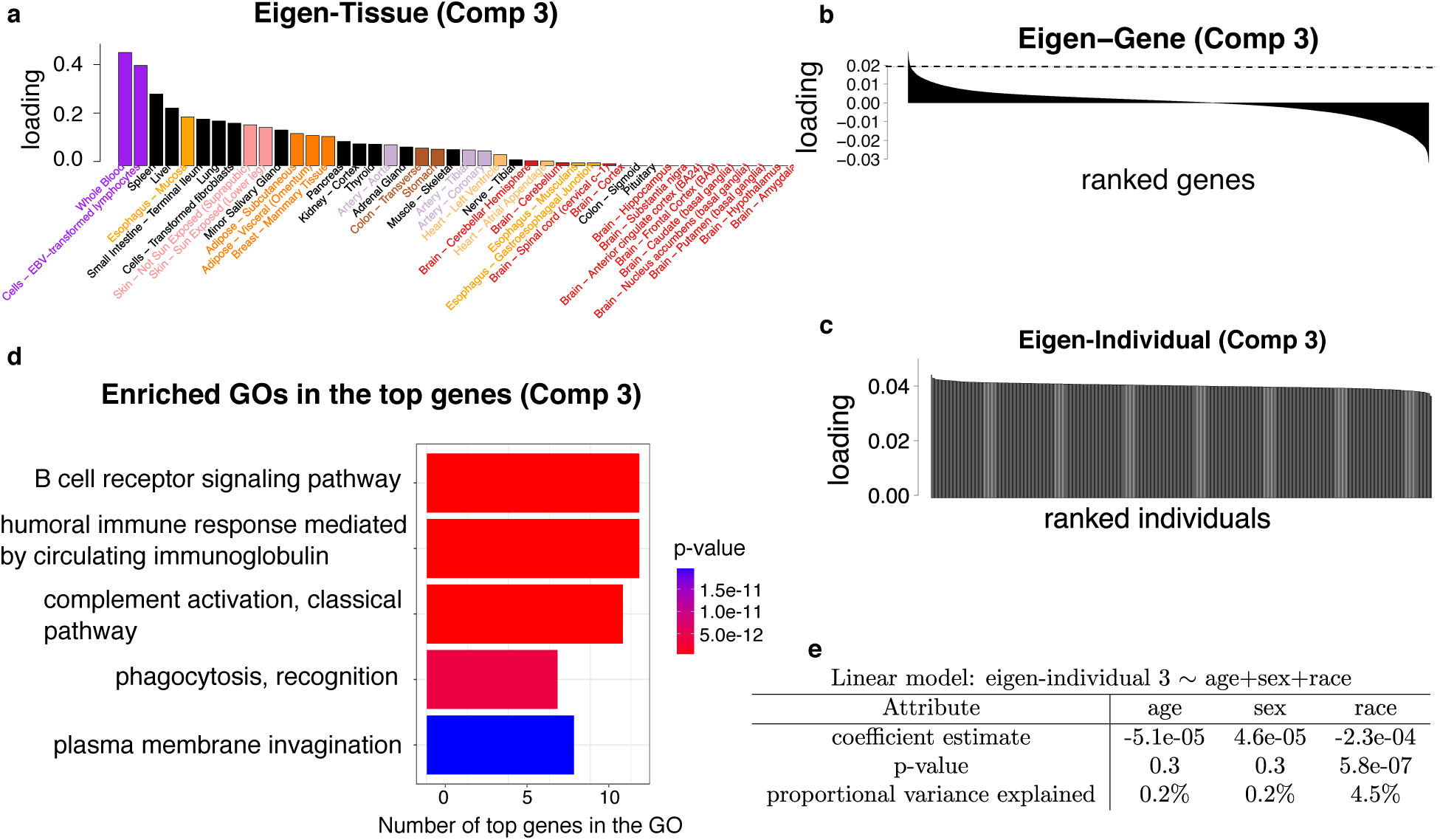
Expression module III - tissues involved in immune response. Panels (a)-(c) represent the triplets of sorted singular vectors, whereas panels (d)-(e) represent the biological contexts of the identified three-way cluster. (a) Barplot of the sorted tissue loading vector, where each tissue is colored based on their functional similarity. (b) Barplot of the sorted gene loading vector, where the dotted line represents the threshold for the top genes. (c) Barplot of the sorted individual loading vector. (d) Enriched GO annotations among the top 89 genes identified from the gene loading vector. The enrichment *p*-values are obtained from hypergeometric tests with B-H correction. The GO size on the axis represents the number of top genes that belong to the GO annotation. (e) Linear regression analysis of individual loadings against individual-level covariates (age, sex and race). The proportional variance explained is computed from ANOVA analysis.

*Component IV - tissues with structural similarities*. The fourth gene expression module is enriched for collagen catabolic process (e.g. *COL8A1, COL15A1, COL3A1*; *p* = 4.5 × 10^−12^), collagen fibril organization (e.g. *DPT, LUM, COL14A1*; *p* = 1.1 × 10^−11^), and multicellular organismal catabolic process (e.g. *MMP2, COL6A2, OL1A1*; *p* = 1.5 × 10^−11^) (Figure 7d). These GO terms represent the synthesis of extracellular matrix proteins that give structure support and elasticity to tissues. For example, the top-ranked gene, *MYOC*, encodes the protein *myocilin*, which is believed to play an important role in cytoskeletal structure (Fishilevich et al. 2016). The second-ranked gene, *PLN*, is associated with the protein *phospholamban*, which regulates the activity of cardiac and smooth muscle (Fishilevich et al. 2016). Coupled with this gene cluster, the tissue cluster is heavily loaded on three artery subtypes (tibial, aorta, coronary). Other highly-ranked tissues include adipose, esophagus, heart, and colon (Figure 7a). These organs are rich in connective fibers with stretch-recoil properties, endowed with the molecular functions we identified earlier. Furthermore, we found that the genes driving this module are expressed at slightly lower rates in women and African-Americans (Figure 7e). The top sex-related gene is XG (X-linked blood group), and the top race-related gene is TUSC5 (Tumor suppressor candidate 5).

**Figure 7.**
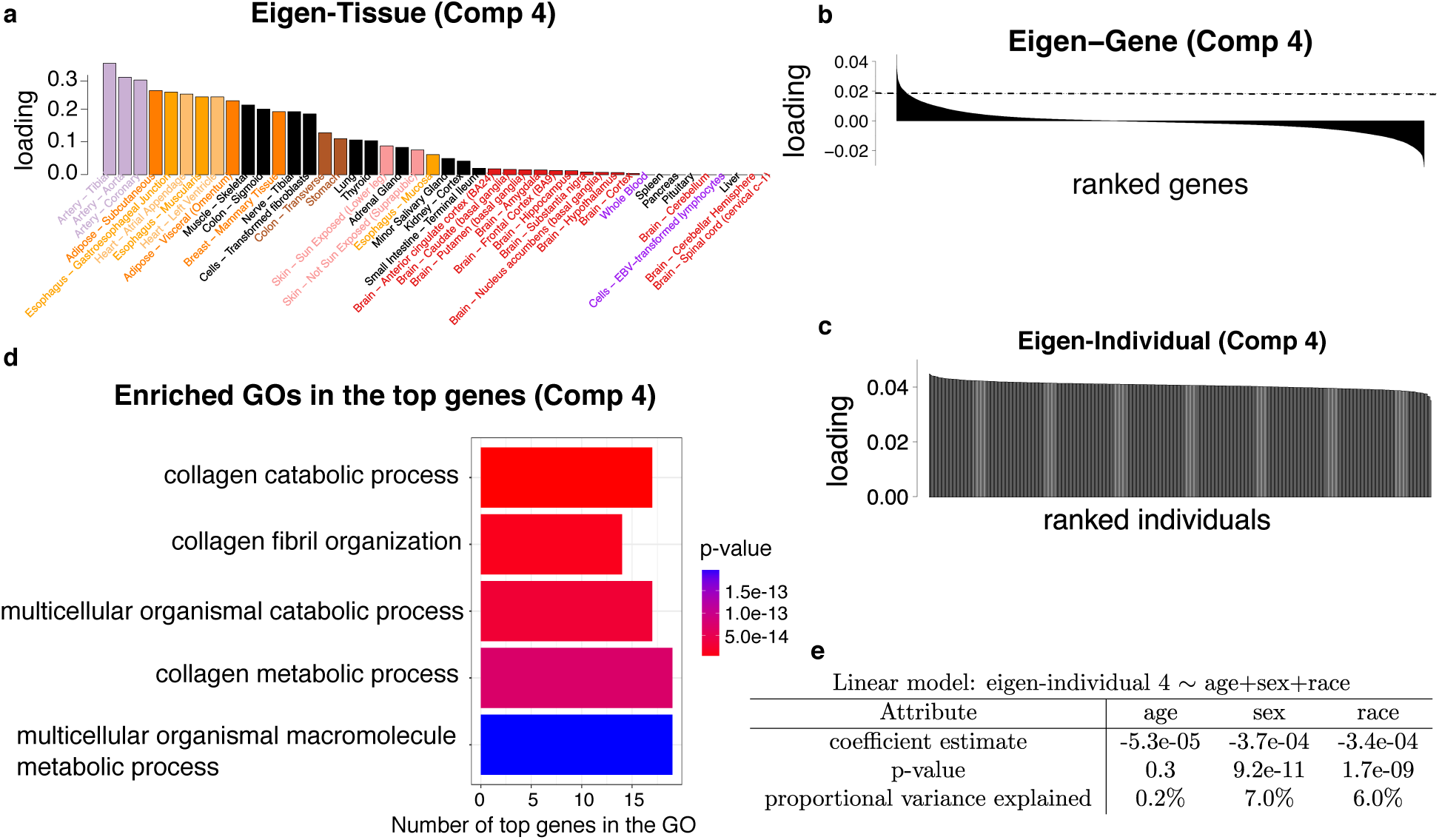
Expression module VI - tissues with structural similarities. Panels (a)-(c) represent the triplets of sorted singular vectors, whereas panels (d)-(e) represent the biological contexts of the identified three-way cluster. (a) Barplot of the sorted tissue loading vector, where each tissue is colored based on their functional similarity. (b) Barplot of the sorted gene loading vector, where the dotted line represents the threshold for top genes. (c) Barplot of the sorted individual loading vector. (d) Enriched GO annotations among the top 352 genes identified from the gene loading vector. The enrichment *p*-values are obtained from hypergeometric tests with B-H correction. The GO size on the axis represents the number of top genes that belong to the GO annotation. (e) Linear regression analysis of individual loadings against individual-level covariates (age, sex and race). The proportional variance is computed from ANOVA analysis.

*Other expression modules identified in the global analysis*. Each of the remaining expression modules is active in only a subset of tissues, indicating the presence of tissue specificity (Supplemental Table S1). These detected modules are specific to skin (exposed and non-exposed), cell lines (EBV-transformed lymphocytes and transformed fibroblasts), liver, muscle (skeletal and cardiac), and cerebellar regions (Supplemental Table S1). Of note is the strong signal of underexpression in the cerebellum in women relative to men. We also found that tissues derived from the same embryologic origin tend to be clustered together; for example, the 4th eigen-tissue groups together blood vessels, heart, smooth muscles and connective tissues (mesoderm), whereas the 7th eigen-tissue groups together liver, pancreas and kidney (endoderm). As seen in Supplemental Figure S2-Supplemental Figure S7, the skin-specific module is enriched with keratin-related genes, the two cell lines are enriched with genes responsible for cell division (e.g. chromosome segregation, meiosis, sister chromatid segregation), the liver-specific module is enriched with inflammation and triglyceride related genes, the muscle-specific module is enriched with myofibril related genes, and the cerebellum-specific gene module is enriched for excretion and dorsal spinal cord development. All of these ontologies are consistent with the function of the associated tissues. Conversely, most eigen-individuals have limited descriptive power compared to the eigen-gene and eigen-tissue (Supplemental Table S1). This was expected because variation in gene expression is usually much lower among individuals than among tissues (Melé et al. 2015). In a subsequent section (Fine structures in subtensors of similar tissues), we detail how targeted analyses of subtensors comprised of similar tissues help mitigate this effect by revealing patterns of variation which are overwhelmed by the large heterogeneity in tissues at the global level.

*Relationship between different expression modules*. The aforementioned modules are naturally ranked according to the amount of variability explained by the associated tensor components (Materials and Methods). We chose to focus on the top 10 components based on their interpretability. Unlike matrix SVD, the components obtained from tensor decomposition are usually not guaranteed to be orthogonal (Supplemental Figure S8). Examination of pair-wise correlations revealed relatively low levels of overlap between most eigenvectors. We did find modest agreement (Pearson’s *r* = 0.44) between gene cluster 4 (tissues with smooth and cardiac muscle) and gene cluster 9 (skeletal and cardiac muscle) (Supplemental Figure S8). Given that these clusters both correspond to muscle-driven eigen-tissues, the similarity is sensible. Likewise, tissue cluster 8 represents a mixture of blood and muscle tissues and overlaps most with the tissue cluster 3 (blood cluster, *r* = 0.59) and tissue cluster 9 (muscle cluster, *r* = 0.43). Nevertheless, we found that the various components in our GTEx analysis tend to capture non-redundant information (Supplemental Table S1), as later components are able to uncover fine-scale structure nested in earlier components. For example, the two cerebellum tissues are separated from other brain regions in the 10th component (Supplemental Figure S7), although they all belong to the larger brain cluster in the 2nd component (Figure 5a). In fact, the eigen-genes in these two clusters prioritize different sets of genes; the brain cluster prioritizes general neural genes such as *OPALIN* and *GFAP*, whereas the cerebellum cluster prioritizes more specific genes such as *ZP2* and *SLC22A31*. Recent study shows that the expression of ZP2 in brain is both spatially-restricted and time-regulated (i.e. transiently enriched during a narrow time window) (Kang et al. 2011), and our analysis demonstrates that ZP2 is almost exclusively expressed in the cerebellum but not in other tissues, including non-cerebellar regions of the brain.

### Fine structures in subtensors of similar tissues

Although our global analysis successfully uncovers distinctive expression patterns in the GTEx data, it may miss finer-scale structure within similar tissues or within similar individuals because of the high degree of inter-tissue heterogeneity. We suspected that restricting the analysis to relatively homogeneous tissues would increase the discriminative power of eigen-individuals and provide information about which genes separate similar tissues. In order to reveal the crucial individual × tissue specificity, we considered six tissue groups, each consisting of morphologically and functionally similar tissues (Materials and Methods). For each tissue group, we built a subtensor and applied *MultiCluster* to identify gene × individual × tissue modules in the subtensor.

*Spatially-restricted and sex/age-related expression in the brain*. Table 1 shows the expression modules detected in the brain subtensor. We found that most expression modules are spatially restricted to specific brain regions, such as the two cerebellum tissues (component 2), three cortex tissues (component 4), and three basal ganglia tissues (component 5). The corresponding gene clusters capture distinctly-expressed genes that are over- or under-expressed in each region. Genes overexpressed in the cerebellum region are strongly enriched for dorsal spinal cord regulation (*p* = 9.8 × 10^−7^) whereas the under-expressed genes are most strongly enriched for forebrain development (*p* = 3.4 × 10^−8^). An opposite enrichment pattern is observed for basal ganglia region. This is consistent with the spatial location of the cerebellum (located in the hindbrain) and basal ganglia (which is situated at the base of the forebrain). In addition, we noticed an abundance of over-expressed *HOX* genes in the spinal cord (cervical C-1) compared to other brain regions (Figure 8a). The *HOX* gene family (*HOXA–HOXD*) is a group of related genes that control the body plan and orientation of an embryo. The non-uniform expression of *HOX* genes across brain regions may suggest the particularly important role of the spinal cord during early embryogenesis.

**Table 1.**
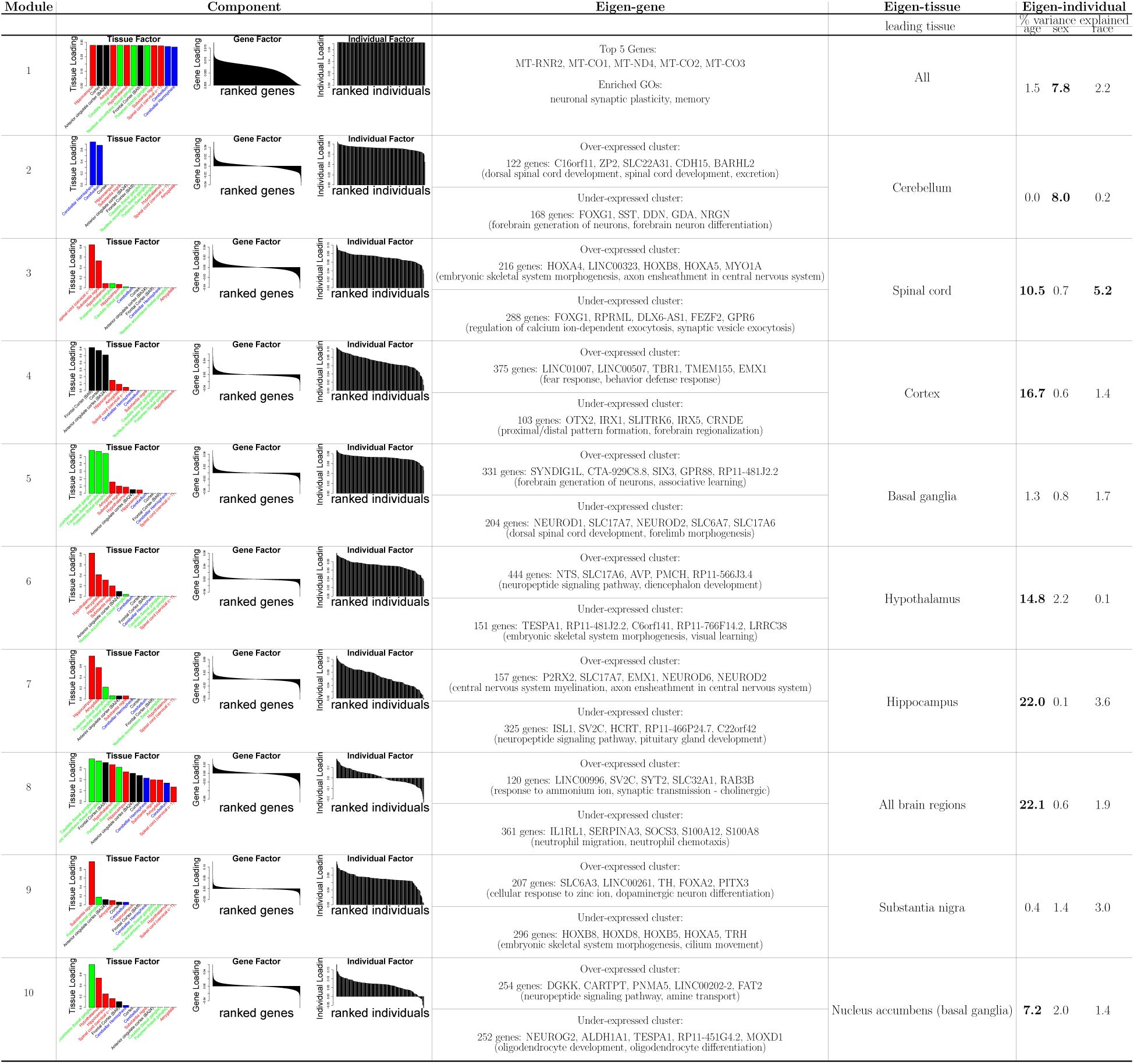
Top 10 expression modules in the brain subtensor. The brain subtensor contains two cerebellar tissues (cerebellar hemisphere, cerebellum; coded in blue), three basal ganglia tissues (caudate, nucleus accumbens, and putamen; coded in green), three cortex tissues (cortex, anterior cingulate cortex-BA24, frontal cortex-BA9; coded in black) and other four tissues (spinal cord-cervical c-1, substantia nigra, hypothalamus, hippocampus; coded in red). The top 10 expression modules (ranked by their singular values) were identified from the *MultiCluster* method. For each component, we plotted the barplots for the sorted tissue loadings, gene loadings, and individual loadings, respectively. In each eigen-gene, we listed top/bottom 5 genes as well as the enriched GO annotations in the identified clusters. In each eigen-tissue, we reported the leading tissue with the largest tissue loading. In each eigen-individual, we reported the proportional variance of the individual loadings that was explained by age, sex, or race, respectively. Number in bold indicates *p* < 10^−3^.

**Figure 8.**
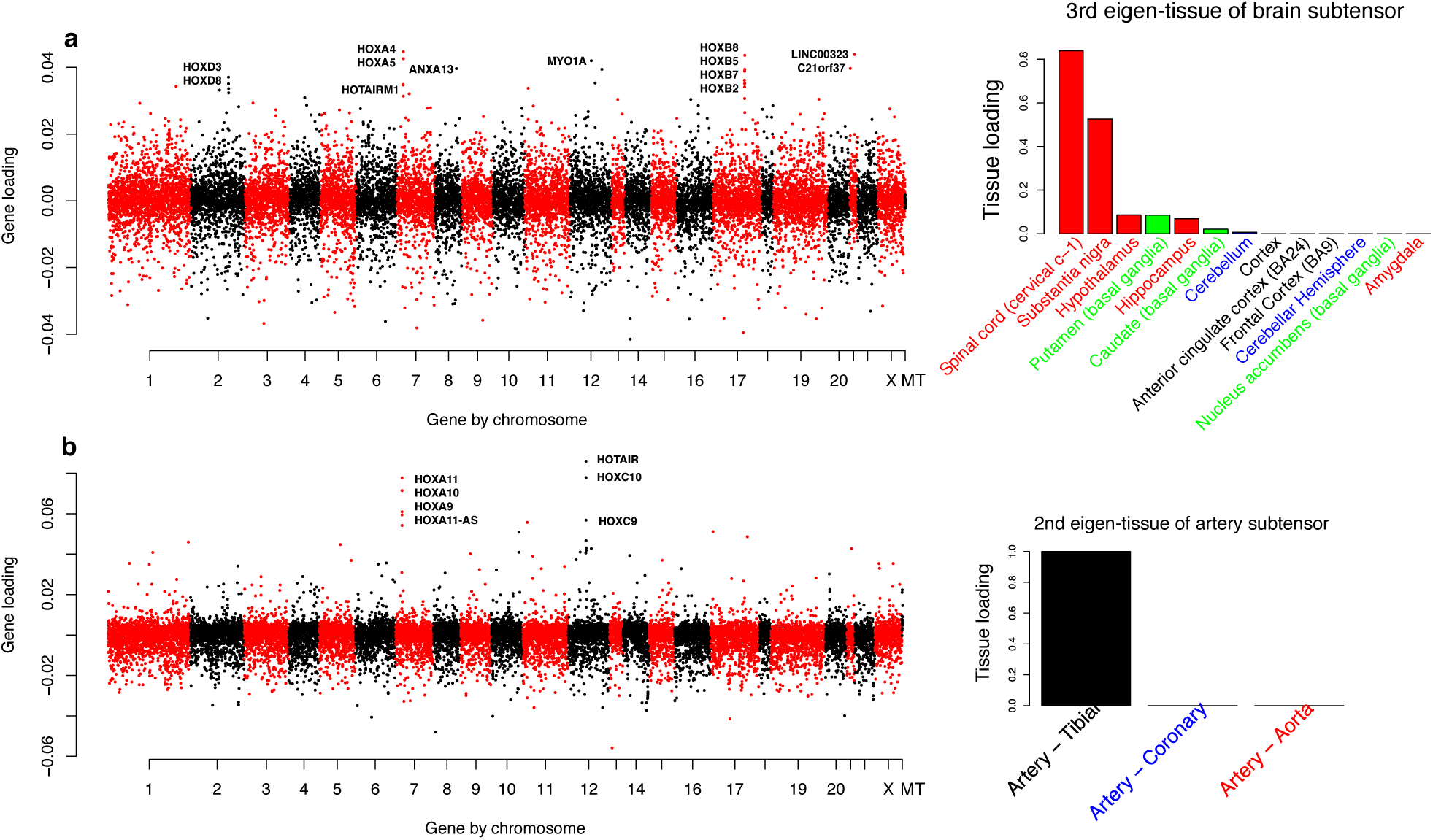
HOX gene expressions and associated tissue loadings in different subtensors. (a) Over-expression of *HOX* genes in spinal cord (cervical C-1) compared to other brain tissues. This expression pattern was identified from the 3rd tensor component of the brain subtensor. (b) Over-expression of *HOX* genes in tibial tissues compared to other artery tissues. This expression pattern was identified from the 2nd tensor component of the artery subtensor. In each panel, the left figure plots gene loadings against gene positions on the chromosomes. Genes with extreme loadings (e.g., *HOXD* genes, *HOXB* genes, *HOXA* genes, etc) are labeled on the plot. The right figure shows the barplot of tissue loadings in the corresponding eigen-tissue.

In addition to tissue-specificity, most expression modules also exhibit considerable individual-specificity. We identified two sex-related and six age-related expression modules from the top tensor components (bolded in Table 1). The second gene module is found to be both cerebellum-specific and sex-related. By ranking genes based on their *p*-values for sex effect in the direction of eigen-tissue (Materials and Methods), we found that the top sex-related gene in this module is X-Y gene pair *PCDH11X/Y*. In fact, the combined expression of *PCDH11X/Y* is significantly lower in the cerebellum (paired t-test *p*-value < 2 × 10^−16^) and in females (*p* = 8.0 × 10^−11^), and the expression level also decreases with age (*p* = 3 × 10^−3^). Notably, *PCDH11X* was the first reported gender-linked susceptibility gene for late-onset Alzheimer’s disease (Carrasquillo et al. 2009), and it may also be implicated in developmental dyslexia (Veerappa et al. 2013). However, its homologous gene on the Y-chromosome, *PCDH11Y*, is believed to have different expression regulation. Studies show that *PCDH11X* and *PCDH11Y* are differentially regulated by retinoic acid. This acid stimulates the activity of *PCDH11Y* but suppresses *PCDH11X* (Priddle and Crow 2013), perhaps explaining the sex-specificity we observed for this gene pair in most brain tissues.

Significant age effects are widely present in the identified expression modules (Table 1). In particular, age explains over 15% individual-level variation in module 4 (cortex), module 7 (hippocampus), and module 8 (all brain tissues). Notably, the hippocampus is associated with memory, in particular long-term memory, and is vulnerable to Alzheimer’s disease (Lam et al. 2017). In the module 4, *GPR26* is found to be the top age-related gene. Using linear regression models (Materials and Methods), we confirmed the significant decrease of GPR26 expression with age in all three cortex tissues (cortex, *p* = 1.9 × 10^−18^; frontal cortex, *p* = 8.8 × 10^−12^, anterior cingulate cortex, *p* = 1.9 × 10^−7^) but not in the substantia nigra (*p* = 0.17) or cerebellum (*p* = 0.64). It is worth noting that both the substantia nigra and cerebellum have zero loadings in the eigen-tissue (Table 1), so our tensor-based approach automatically detects the tissue-specificity of this aging pattern. In line with our findings, a recent study shows that *GPR26* plays an important role in the degradation of intranuclear inclusions in several age-related neurodegenerative diseases (Mori et al. 2016).

Another age-related module is component 8 (Table 1; age explains 22.1% individual-level variation), in which every tissue has a non-zero loading in the eigen-tissue. We found that the 2nd top gene in this expression module is *SERPINA3*, a gene implicated in Alzheimer’s disease (Ciryam et al. 2016). This gene has a moderate age effect in each single tissue (*p*-values ranging from 1.1 × 10^−6^ to 0.05 with all effects in the same direction). Using tensor projection, we obtain *p* = 1.0 × 10^−8^ in the direction of eigen-tissue, improving the single-tissue analysis by two orders of magnitude. This increased power is one example of an advantage our tensor-based approach via the sharing information across similar tissues. See Supplementary Data for the complete list of *p*-values for covariate-associated genes obtained using our approach.

*Tissue-specific and race/sex-related expression in cardiac and skeletal muscles*. The muscle subtensor consists of gene expression profiles sampled from two heart regions of cardiac muscle (atrial appendage, left ventricle) and skeletal muscle. As seen from Supplemental Table S2, the top five eigen-tissues reveal the hierarchy-based similarity among the three tissues. Eigen-tissues 2 and 3 represent the muscle and heart clades, respectively, whereas eigen-tissues 4 and 5 further partition the heart clade into each of its constituent components. The corresponding gene clusters capture the differentially expressed genes which drive the tissue partition. In particular, the 3rd gene cluster comprises 511 genes that are similarly expressed in the heart tissues but have distinctive expression patterns in the heart relative to skeletal muscle. By projecting the expression tensor through the 3rd eigen-tissue (Materials and Methods), we identified 122 race-related genes and 95 sex-related genes in this gene cluster (Supplementary Data). Comparatively, the corresponding single-tissue analyses uncover only 91 race-related (86 sex-related) in the left ventricle and 107 race-related (91 sex-related) genes in the atrial appendage, again displaying the increased detection power of our tensor method for genes with moderate but concordant effects across tissues. One such covariate-associated gene, *TCF21*, is a tumor suppressor gene involved in dilated cardiomyopathy which exhibits consistently decreased expression in African-Americans (*p* = 4.7 × 10^−12^ in the ventricle; *p* = 8.1 × 10^−17^ in the atrial appendage), in females (*p* = 1.8 × 10^−6^ in the ventricle; *p* = 2.3 × 10^−12^ in the atrial appendage), and in the elderly (*p* = 9.4 × 10^−4^ in the ventricle; *p* = 2.8 × 10^−9^ in the atrial appendage). Using our tensor-based joint analysis to test for individual effects, we improve the statistical significance to *p* = 2.9 × 10^−20^ for race effect, 3.3 × 10^−13^ for gender effect, and 1.3 × 10^−9^ for age effect, respectively.

*Gender-driven distinction between breast and two adipose tissues*. In the adipose subtensor, we detected several modules representing highly expressed genes in breast tissue and females. For example, the eigen-tissue in module 2 (Supplemental Table S3) loads on breast tissue only and 40% of the individual-level variation is attributable to sex. Such a pattern highlights the gender-driven distinction between breast tissue and the other two adipose (subcutaneous and visceral) tissues. More importantly, the top five genes in this module (*SCGB2A2, KRT17, VTCN1, PIP, MUCL1*) are breast cancer biomarkers (Barh 2014; Lacroix 2006; Merkin et al. 2017; Naderi and Vanneste 2014). Each of these five genes has a significant sex effect in the breast tissue (*p* ranging from 9.8 × 10^−8^ to 1.1 × 10^−18^), but not in the other two adipose tissues (*p* ranging from 0.003 to 0.62). In addition to the risk genes themselves, genes known to be co-expressed with them also tend to be included in this module. Indeed, the secretoglobins *SCGB1D2* and *SCGB2A1*, known to be reliably co-expressed with *SCGB2A2* (Lacroix 2006), were also highly ranked (13rd and 51st, respectively) in the gene cluster.

*Tissue-specific and age-related expression in three artery types*. In the artery subtensor, the most distinct tissue is the tibial artery, with the 2nd eigen-tissue clearly separating it from the other two arteries (coronary and aorta). The corresponding eigen-gene peaks in the *HOXA* and *HOXC* regions (Figure 8b), indicating the overexpression of *HOX* genes in the tibial artery relative to the coronary artery and aorta. Of note is the famous lncRNA *HOTAIR*, the first RNA gene found to regulate distantly located genes throughout the genome. *HOTAIR* gene is located inside the *HOXC* locus and plays a key role in the initiation and progression of different types of cancer. In addition, this tibial-specific expression module exhibits significant age-relatedness; in particular, 14.9% of the individual-level variation is attributable to age. A further investigation reveals that this aging signal is mostly driven by the group of genes at the negative end of the eigen-gene (Supplemental Table S4). Among the 517 genes in the cluster, we detected 207 age-related genes (with significance threshold α = 10^−3^/517 ≈ 1.9 × 10^−6^ via Bonferroni correction), 206 of which are over-expressed with age (Supplementary Data). The top age-related gene is ARHGEF28, encoding a member of the Rho guanine nucleotide exchange factor family. The encoded protein may be involved in amyotrophic lateral sclerosis (ALS), a neurodegenerative disorder that affects the movement of arms, legs, and body (Droppelmann et al. 2013).

*Expression patterns in reproductive tissues*. The subtensor analyses of gender-specific reproductive tissues (ovary, uterus, and vagina for female; prostate and testis for male) also reveal interesting gene expression patterns. We observed a clear uterus × age specificity in the female subtensor (Supplemental Table S5) and a prostate × age/race specificity in the male subtensor (Supplemental Table S6).

In the female tensor, component 4 is an age-related expression module which also distinguishes the uterus from the other two tissues (ovary, vagina). The top genes in this gene cluster are *CHRDL2* (also known as *BNF1*, Breast Tumor Novel Factor 1), *DPP6* (Dipeptidyl Peptidase VI), *TEX15* (Testis Expression 15) and *ZCCHC12* (Zinc Finger CCHC-Type Containing 12). These genes tend to be related to reproductive functions such as DNA double-strand break repair (*TEX15*), or be involved in X-linked disease (*ZCCHC12*). DPP6 expression changes are associated with age and is preferably expressed in the uterus compared to the other two tissues (paired t-test *p* < 2 × 10^−16^). In particular, *DPP6* expression decreases significantly with age in the uterus only (*p*-value for age = 1.3 × 10^−16^ compared to *p* = 0.05 in ovary and *p* = 0.51 in vagina). A recent study (Chettier et al. 2014) reveals that *DPP6* harbors a copy number variant locus, rs758316, that is significantly associated with endometriosis. While the function of *DPP6* in uterus remains unclear, its unique aging pattern makes it a good candidate for further investigation.

In the male subtensor, we found that prostate and testis are characterized by two distinct clusters of genes (components 2 and 3 in Supplemental Table S6). Genes over-expressed in prostate are mostly related to prostate glandular acinus development (e.g. *HOXB13*, *FOXA1*) and fluid transport (e.g. *SLC14A1*, *UPK3A*). Among the top five genes, *KLK3, ACPP*, and *MSMB* encode the three predominant proteins secreted by a normal human prostate gland. Their protein level in serum is commonly used for monitoring prostate disorders such as prostatitis (*KLK3* and *MSMB*) or prostate cancer (*ACPP, MSMB, HOXB13*). Genes over-expressed in testis, on the other hand, are mostly enriched for sperm motility (e.g. *TNP1, AKAP4, SMCP*; *p* = 1.4 × 10^−17^), meiosis I (e.g. *DMRTC2, SYCP1, BRDT*; *p* = 7.9 × 10^-17^), and male meiosis (e.g. *BRDT, DDX4, TEX15*; *p* = 5.1 × 10^−7^). In addition, we found that the prostate-specific modules tend to be race- or age-related (components 2, 4, 7, 8 and 9 in Table S6). The top race-related gene in module 2 is *SPINK2*, with a higher average expression in black than white Americans (*p* = 9.0 × 10^−4^). *SPINK2* encodes a serine protease inhibitor located in the spermatozoa, and recent evidence shows that *SPINK2* deficiency leads to fertility changes by causing sperm defects in individuals with one defective copy and azoospermia in those with two defective copies (Kherraf et al. 2017). Given the high degree of race-relatedness among gene expression patterns in the prostate, it is relevant to note the large discrepancy between prostate cancer rates in black and white Americans (153.9 vs 86.8 per 100,000, respectively, according to U.S. Cancer Statistics Working Group (2017)). Although the GTEx cohort does not include individuals with cancer, the strong dependence of prostate cancer incidence rates on race suggests that some of the genes identified as race-related may be involved in the development of prostate cancer and merit further study.

### Common expression features in subtensors

Analysis of subtensors allows us to focus on one tissue group at a time, revealing a finer-scale characterization of transcriptional variation in different parts of the body. In addition to tissue comparisons within each subtensor, it is interesting to examine how expression modules identified in different tissue groups compare, and several intriguing features emerge from this meta-analysis.

We found that genes belonging to the same family tend to be clustered closely together. For example, the genes *ZIC1* and *ZIC4* always co-occur in gene clusters (Supplemental Figure S9, component 3 in Supplemental Table S3, component 6 in Supplemental Table S1). *KRT13* is often paired with *KRT4* (somponent 10 in Supplemental Table S2 and Supplemental Table S4), and *HBA1/HBA2* have similar tissue loadings (Supplemental Figure S10). As we reduce the dimension of expression data from thousands genes to a handful of eigen-genes, these co-expressed genes provide a validation of our gene groups. Many gender-related expression modules also prioritize *XIST* (e.g. component 5 in Supplemental Table S4 and components 5, 6, 8 in Supplemental Table S3). In fact, *XIST* is one of the most famous lncRNA genes essential for X-inactivation process and female survival (da Rocha and Heard 2017. The wide presence of *XIST* in our analysis reinforces its crucial role in gender-differentiated expression in tissues.

Several subtensors yield similar eigen-genes, suggesting the presence of common expression patterns shared by seemingly unrelated tissues. In particular, we identified three eigen-genes, one in each of the female and male subtensors and another in the artery subtensor (Supplemental Figure S11) that exhibit clearly similar gene loadings. The three eigen-genes are mainly loaded in four genomic loci encoding immunoglobulin: the *IGK, IGJ, IGH*, and *IGL* regions on chromosomes 2, 4, 14, and 22, respectively. Among other top genes, we identified *FCRL5, CH3L1*, and *CHIT1*, all of which are related to immune response. The repeated appearance of this expression module in different tissues and individuals highlights the similar roles of distinct tissues in disparate bodily systems. Despite its presence in each of these tissue groups, this module exhibits differential expression between tissues within each tissue group. Namely, these immune genes are more expressed in the vagina, prostate, and aorta compared to the other reproductive tissues (ovary/uterus, testis) (Supplemental Figure S11) and artery types (tibial/coronary). Interestingly, this module exhibits a strong race effect in the prostate, with higher average expression in black than white men (explaining 14% variation in the corresponding eigen-individual, *p* = 3.1 × 10^−9^; see component 7 in Supplemental Table S6). Such non-uniform expression reflects the complex relationship between related tissues and individuals which our method is well suited to uncover.

## Discussion

We have presented a new multi-way clustering method, *MultiCluster*, and demonstrated its utility in identifying three-way gene expression patterns in multi-tissue multi-individual experiments. To the best of our knowledge, our model is the first tensor decomposition that uses nonnegativity constraints and the sharing of information across modes to detect multi-modal specificities in this context. We are able to highlight gene expression modules that are common to all tissues/individuals or exclusive to particular tissue × individual combinations. Our method uncovers three-way specificities with clear statistical and biological significance in both simulations and the GTEx dataset. We have provided evidence that the distinctions among human tissue gene expression profiles are usually driven by a small set of functionally coherent genes and that many age-, race-, gender-related genes exhibit tissue-specificity even within functionally similar tissues.

A major benefit of *MultiCluster* (and tensor-based methods in general) over existing tissue comparison method (Consortium et al. 2015) is the substantially reduced number of comparisons which must be considered. If one wanted to analyze every possible tissue pairing in a set of *n* tissues, roughly *n^2^* analyses would have to be performed and the results would need to be synthesized via a meta-analysis. The analysis can be even more prohibitive if one wanted to examine the 2*^n^* possible tissue-specificity and sharing configurations (Consortium et al. 2015). In contrast, *MultiCluster* identifies coherent clusters across each mode of the data in a single step and associates the resulting variation with biological contexts. Though prior knowledge of tissue function can greatly reduce the number of pairwise comparisons, doing so constrains potential insights to the set of hypothesized tissue modules. For instance, components III and IV of our global tensor decomposition consist of diverse tissues which may not have been grouped together a priori. An additional benefit of *MultiCluster* is the ranking of tissue modules by amounts of variation.

We also implemented a tensor projection procedure to test whether differentially-expressed genes correlate with biological attributes (age, sex or race) and found that we achieve improved power relative to single-tissue tests. This approach can be naturally extended to (trans-)eQTL analyses by testing the projected expression of each gene against genetic variants across the genome. Alternatively, one can test each individual loading vector against genetic variants to identify eQTLs (Hore et al. 2016). Existing multi-tissue eQTL analyses usually proceed by identifying eQTLs in each tissue separately before combining single-tissue results via meta analysis (Battle et al. 2017). However, the large numbers of genes, tissues, and genetic variants potentially incur a substantial penalty for multiple testing and there is also the risk of under-powered tests due to limited sample sizes. An avenue worthy of pursuit is to apply *MultiCluster* for eQTL discovery in large multi-tissue expression studies.

Several consortium efforts — including the Cancer Genome Atlas (Weinstein et al. 2013), the Allen Human Brain Atlas (Hawrylycz et al. 2012), and the Encyclopedia of DNA Elements (Consortium et al. 2004) — have been recently completed or continue to collect data, and our method provides a powerful computational tool to analyze the genome-scale transcriptional profiles that they produce. Although we have presented *MultiCluster* in the context of multi-tissue multi-individual gene expression data, the general framework applies to more general multi-way datasets. One possible extension is integrative analysis of omics data (Berger et al. 2013), in which multiple types of omics measurements (such as gene expression, DNA methylation, microRNA) are collected in the same set of individuals (Weinstein et al. 2013). In such cases, tensor decomposition may be applied to a stack of data matrices or correlation matrices, depending on the specific goals of the project. Other applications include multi-tissue gene expression studies under different experimental conditions in which one may be interested in identifying 4-way expression modules arising from the interactions among individuals, genes, tissues, and conditions. The tensor framework can also be applied to time-course multi-tissue gene expression (Almon et al. 2003). In this instance one may treat time as the 4th mode and extend the tensor projection approach to identify the time trajectories of 3-way expression modules.

One assumption made by our algorithm is that expression matrices for different tissues are of the same dimension. In the present work, we obtain a complete tensor prior to decomposition by adopting a robust imputation scheme. A possible approach which avoids the need for imputation is to make use of the connection between tensor decomposition and joint matrix factorization (Hore et al. 2016; Xiao et al. 2014; Kuleshov et al. 2015). For example, one could model the *n_G_*-by-*n_It_* expression matrix **M**_*t*_, where t indexes the tissue, as ***M**_t_* ≈ ***AΛ**_t_**B**_t_* with some identifiability conditions. This model is a relaxation of tensor decomposition because it allows different tissues to have different column (individual) spaces ***B**_t_* while sharing the same row (gene) space ***A***. The diagonal matrix ***Λ**_t_* captures the tissue-sharing and specificity as before. Another potential approach is to implement tensor imputation and decomposition simultaneously via a low-rank approximation, an idea which has roots in the matrix literature (Troyanskaya et al. 2001; Candès and Recht 2009). We did not pursue this direction beyond initial investigation because of the computational cost and bias introduced by the non-random sample collection in the GTEx data.

## Materials and Methods

### Data Processing

Here we describe our data processing steps.

#### Normalization and quality control

To prepare for comparisons across samples, normalization was performed using the size factors produced by the *estimateSizeFactors* function of DESeq2 (Love et al. 2014). After normalizing, we applied quality control measures at both the tissue and gene levels to refine our results and restrict our analyses to informative features. Specifically, we required at least 15 samples to include a given tissue and an average of at least 500 normalized reads in one or more tissues to retain a gene.

#### Correction for nuisance variation

There were several technical covariates whose effects we wished to remove in order to focus on the correlation between gene expression and biological and phenotypic characteristics. The choice of these factors was driven by a preliminary step in which we looked for signs of significant correlations between any one technical covariate and expression levels. After curating the list of technical covariates in this manner, we were left with the sample collection cohort (postmortem, organ donor, surgical), ischemic time (IT, in minutes), whether the patient died while on a ventilator, and the date of RNA sequencing. Evidence of effects due to some of these factors has been discussed previously elsewhere (McCall et al. 2016).

To correct for the variation due to these factors while preserving the impact of phenotypes, we ran multiple linear regression for every tissue-gene pair per the following linear model:

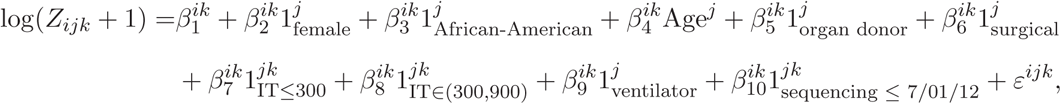

where 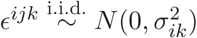. Here *Z_ijk_* is the normalized read count in gene *i*, individual *j*, and tissue *k*. The superscripts on coefficients and covariates indicate to which attribute(s) (gene, individual, tissue) they correspond.

After fitting this set of models, we removed the estimated effects due to the aforementioned technical covariates. To obtain the log-transformed corrected expression value *Y_ijk_*, we computed

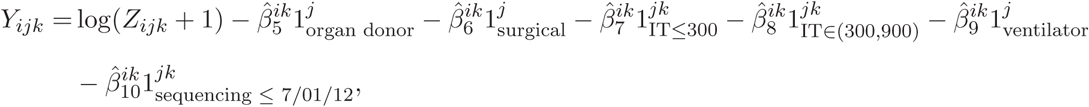

for all *i* = 1,…, *n_G_*, *j* = 1,…, *n_I_*, and *k* = 1,…, *n_T_*, where *n_G_*, *n_I_*, and *n_T_* denote the number of genes, individuals, and tissues, respectively.

#### Imputation of unobserved entries

Applying our tensor decomposition method necessitates a complete set of observations in which we have the RNA-seq gene read counts for all individuals in all considered tissues. To obtain the requisite data structure from the initial incomplete set of observations, we implemented a *k*-nearest neighbors imputation scheme which fills missing entries with the averaged read counts from the corresponding tissue in the ten individuals most similar in terms of age, race, and gender. This method preserves the pre-imputation signal in the data and does not appear to introduce erroneous clusterings due to the non-random sample collection procedure as validated by comparing hierarchical tissue trees (Supplemental Figure S12) before and after imputation. For tissue hierarchy, we took the mean across individuals to produce a gene-by-tissue matrix and computed the distance matrix based on the tissue-tissue Spearman correlation matrix. The hierarchy tree was constructed using UPGMA (Unweighted Pair Group Method with Arithmetic Mean) algorithm (Sokal 1958).

#### Handling sex-specificity at the tissue and gene levels

As the GTEx cohort comprises both male and female samples, it does not make sense to compare some tissues and genes when using the full set of individuals. To remedy these concerns, we held out sex-specific tissues (e.g. testis, uterus, etc) and only considered them in smaller analyses in the appropriate gender. Further, we also removed all Y chromosome genes save those in the pseudoautosomal region, whose reads we combine with their X chromosome paralogs.

#### Tissue groups considered in subtensor analysis

The following six tissue groups are considered in the subtensor analysis: (i) 13 brain tissues; (ii) three artery tissues (tibial, aorta, coronary); (iii) two adipose tissues (subcutaneous, visceral) and breast - mammary tissue; (iv) three muscle tissues (heart - atrial appendage, heart - left ventricle, and muscle - skeletal); (v) three female-specific tissues (ovary, uterus, vagina); and (vi) two male-specific tissues (testis, prostate).

### *MultiCluster* via semi-nonnegative tensor decomposition

In a multi-tissue gene expression experiment, the data take the form of an order-3 tensor, *Y* = 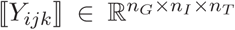 where *Y_ijk_* denotes the (corrected or imputed) expression value of gene *i* measured in individual *j* and tissue *k*. We propose to model the expression tensor *Y* as a perturbed rank-*R* tensor,

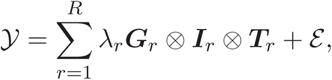

where λ_1_ ≥ λ_2_ ≥ … ≥ λ_R_ ≥ 0 are the singular values in descending order, and *G_r_*, *I_r_*, and *T_r_* are norm-1 singular vectors in ℝ*^n_G_^*, ℝ*^n_I_^*, and ℝ*^n_T_^*, respectively, and 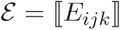 is noise tensor with each entry 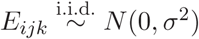. We refer to the vector *G_r_* (respectively, *T_r_* and *I_r_*) as the *r*-th eigen-gene (respectively, eigen-tissue and eigen-individual). We take a successive rank-1 approximation to *Y* (or its residual) by solving the following optimization:

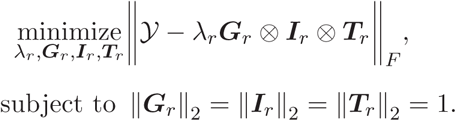

where ‖·‖_F_ is defined entrywise as ‖Y‖ 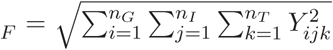 In each iteration, we impose the entrywise non-negativity on the tissue loading vectors *T_r_* ≥ 0 by thresholding negative values of *T_r_* to 0. Then we take the residual tensor *Y* ← *Y* – *λ_r_G_r_* ⊗ *I_r_* ⊗ *T_r_* as the new input and repeat the algorithm to find the next component. The full algorithm is provided in Supplemental Material.

The non-negativity constraint on each *T_r_* eases interpretation of the interaction at the tissue level; a sparse vector *T_r_* implies that the module *r* is “active” in only a few tissues, whereas a dense vector *T_r_* implies that the module *r* is common to several tissues. Without the nonnegativity constraint, it is possible (in fact likely) that each *T_r_* actually contains two expression modules, one which corresponds to the positively-loaded tissues and one to the negatively-loaded tissues. Consequently, gene and individual loadings become less informative because it it difficult to determine with which module they associate.

### Characterizing the expression modules

We applied the *MultiCluster* algorithm to the GTEx tensor with *R* set to 10. Note that *MultiCluster* finds the tensor components via successive rank-1 approximations, so one can always apply the algorithm to the residual tensor if more components are wanted. Here we chose to focus on the top *R* = 10 components based on biological interpretability. For each *r* = 1,…,*R*, we used the procedure discussed below to characterize the biological significance of the loading vectors, *G_r_, I_r_, T_r_*. For ease of presentation, in what follows we drop the subscript *r* and simply write *G, I*, and *T*.

*GO enrichment based on gene loadings*. Let *G* = {1,…,*nG*} denote all genes in the analysis, and *G* = (*G*_1_,…, *G_nG_*)*^T^* be the eigen-gene of interest. We performed GO enrichment analyses in both the top gene cluster *G*_top_ = {*i* ∊ *G*: *G_i_* ≥ *c*_top_ }, and bottom gene cluster *G*_bottom_ = {*i* ∊ *G*: *G_i_* ≤ *c*_bottom_}. Here ctop and cbottom are cut-off values which determine the cluster sizes. For each declared gene cluster, we assessed the overrepresentation of GO terms using the hypergeometric test as implemented in the *enrichGO* function of ClusterProfiler (Yu et al. 2012). The GO annotations were loaded from *org.Hs.eg.db* (Carlson 2017) and only biological processes (BP) terms with at least 10 genes were considered. To account for multiple testing, we report enrichment *p*-values after applying the Benjamini-Hochberg (B-H) correction.

The cut-off values *c*_top_ and *c*_bottom_ control the significance level of the genes in the declared gene clusters. We employ a permutation-based procedure to determine the cut-off values (and thus gene cluster sizes) with a significance level α = 0.005. Specifically, we generated ten null tensors by randomly and independently permuting genes for every individual-tissue pair (*j, k*), i.e.,

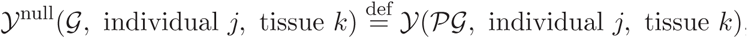

*where PG* represents a random permutation of the set *G* at the pair (*j, k*). We then decomposed each of the null tensors *Y*^null^ and used their eigen-genes *G*^null^ to represent the null distribution of *G*-values. The cut-off value *c*_top_ (respectively, *c_bottom_*) was determined using the top 0.5%-quantile (respectively, bottom 0.5%-quantile) of the empirical distribution of 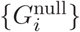
*Effects of biological attributes on individual loadings*. To identify the sources of variation in the individual loadings, we considered the following linear model for an eigen-individual *I* = (1_1_,…, *I_n__i_*)*_T_*,

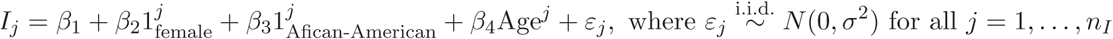

Upon fitting the model, we calculated the proportion of variance explained by each covariate (gender, race, or age) using ANOVA.

### Tensor projection for detecting DE genes

Here we describe our tensor projection procedure for detecting covariate-associated genes. Let *Y* ∊ *R*^*n*_*G*_×*n_I_* ×*n*_*T*_^ denote the expression tensor and *{T_r_* ∊ *R_n_T__* } be the set of eigen-tissues from the decomposition. As we have seen, *T_r_* captures the degree of similarity across tissues in the expression module *r*. Let *Y*(·, ·, *T_r_*) ∊ *R_n_G__*×*n*_*I*_ denote the projection of *Y* through the eigen-tissue *T_r_* = (*T_r_*,1,…, *T_r_, n_T_*)*^T^*; that is, 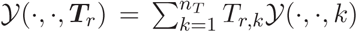. For each gene to be tested, we proposed the following linear model,

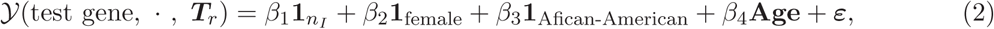

where *ε* ∼ *N*(0, *σ*^2^*I_n_I_×n_I__*), 1*n_I_* denotes a vector of length *n_I_* with every element equal to 1, and *I_n_I__ × *n_I_* denotes an *n_I_*-by-*n_I_* identity matrix. The age-effect was assessed by testing *H*_0_: *β*_4_* = 0 against *H_α_*: *β_4_* ≠ 0. Similar hypothesis testing can be performed for gender and race effects.

In the single-tissue analysis, we considered each gene-tissue pair one at a time and performed the following regression analysis,

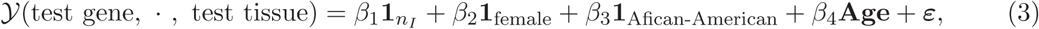

where *ε ∼ N(0, σ^2^I_n_I_×n_I__*), 1*n_I_*. Comparing model (3) with model (2), we see that the tensor projection makes use of the similarity across tissues and thus may boost power for detecting covariate-associated genes.

### Simulation models for assessing three-way clustering

We simulated a set of expression tensors 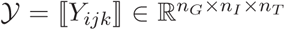, where *n_G_* = 500 (genes), *n_I_* = 500 (individuals), and *n_T_* = 10 (tissues). In each tensor, we created *K_G_* = 5 gene clusters, *K_I_* = 4 individual clusters, and *K_T_* = 3 tissue clusters by randomly assigning each gene (respectively, individual and tissue) to a gene cluster (respectively, individual cluster and tissue cluster) with uniform probability. We generated mean expressions according to *E(Y_ijk_) = μ_1mn_*, where *l, m, n* denote the corresponding gene/individual/tissue cluster that *Y_ijk_* belongs to. We employed three models to simulate the 3-way block means *{μ_1mn_}*.

1. Additive-mean model: 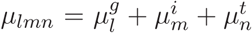, Where 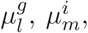, and 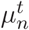 represent the marginal mean for gene cluster *l*, tissue cluster m and individual cluster *n*, respectively. The marginal means, 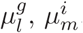, and 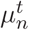, are i.i.d. drawn from N(1,1).
2. Multiplicative-mean model: 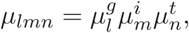where the notation remains the same.
3. Combinatorial-mean model: 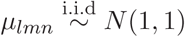; that is, each three-way block has its own mean, independently of each other.

Let *Y*_true_ denote the noiseless tensor with 3-way block means generated from each of the above schemes. The observed expression data were then simulated from *Y* = *Y*_true_ + ɛ, where *ɛ* ∊ ℝ^*n_G_*×*n_I_*×*n_T_*^ is a random Gaussian tensor with each entry i.i.d. drawn from *N(0,σ^2^*).

We simulated 50 expression tensors under each model, and then assessed the accuracy of recovery using the relative error, defined by

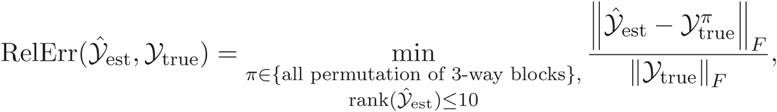

where *ŷ_est_* denotes the rank-*R* approximation (*R* = 1,…, 10) obtained from tensor decomposition.

### Simulation models for detecting DE genes

In order to simulate tensors with both block structure and age signals, we modified the earlier additive model into

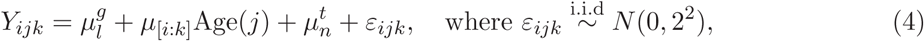

where *Y_ijk_* denotes the expression level of gene *i*, individual *j* and tissue *k*; 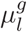 and 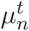 denote the same parameters as before (the marginal means for gene cluster l and for tissue cluster *n*); and

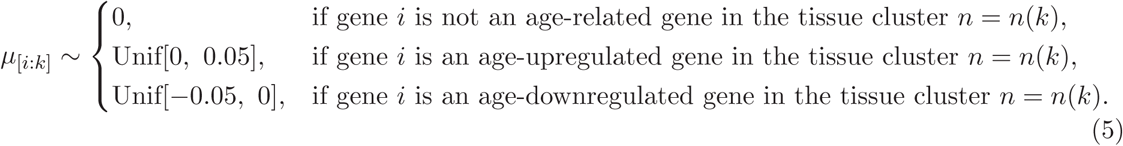

We simulated 50 tensors ℝ^*n_G_*×*n_I_*×*n_T_*^ with *n_G_* = 500 (genes), *n_I_* = 50 (individuals), and *n_T_* = 10 (tissues). In each tensor, we planted five gene clusters, three tissue clusters, and further assigned 100 genes to be age-related (50 up-regulated and 50 down-regulated) in at least one of the three tissue clusters. The ages of individuals were generated using i.i.d. Unif[40, 70], and the effect sizes were simulated using (5). The final expression data was generated based on model (4).

## Runtime

The run times were evaluated using a single processor on an Macbook (Mac OS High Sierra 10.13) with Intel Core i5 2.9GHz CPU and 8GB RAM. We chose the multi-threading option in *SDA* with 4 threads. Both *MultiCluster* and *HOSVD* were implemented in Matlab R2016a 64-bits.

### Data and software availability

Supplementary data containing inferred gene modules and the software *MultiCluster* are available.

## Acknowledgments

We thank Junhyong Kim for helpful discussions and comments on our work. This research is supported in part by a Math+X Research Grant from the Simons Foundation, a Packard Fellowship for Science and Engineering, and a National Institutes of Health grant R01-GM094402. YSS is a Chan Zuckerberg Biohub investigator.

